# Dynamic changes in large-scale functional connectivity prior to stimulation determine performance in a multisensory task

**DOI:** 10.1101/579938

**Authors:** Edgar E. Galindo-Leon, Karl J. Hollensteiner, Florian Pieper, Gerhard Engler, Guido Nolte, Andreas K. Engel

## Abstract

Complex behavior and task execution require fast changes of local activity and functional connectivity in cortical networks at multiple scales. The roles that changes of power and connectivity play during these processes are still not well understood. Here, we study how fluctuations of functional cortical coupling across different brain areas determine performance in an audiovisual, lateralized detection task in the ferret. We hypothesized that dynamic variations in the network’s state determine the animals’ performance. We evaluated these by quantifying changes of local power and of phase coupling across visual, auditory and parietal regions. While power for hit and miss trials showed significant differences only during stimulus and response onset, phase coupling already differed before stimulus onset. An analysis of principal components in coupling at the single-trial level during this period allowed us to reveal the subnetworks that most strongly determined performance. Whereas higher global phase coupling of visual and auditory regions to parietal cortex was predictive of task performance, a second component revealed a reduction in coupling between subnetworks of different sensory modalities, probably to allow a better detection of the unimodal signals. Furthermore, we observed that long-range coupling became more predominant during the task period compared to the pre-stimulus baseline. Taken together, our results show that fluctuations in the network state, as reflected in large-scale coupling, are key determinants of the animals’ behavior.

## Introduction

The brain continuously integrates information from different sensory systems enhancing its ability to filter noisy signals, to construct coherent percepts and guide appropriate actions ^1–4^. Such processes require the flexible and coordinated orchestration of distributed neuronal populations within and across brain areas ^5–9^. Numerous studies have shown that this orchestration relies mostly on three modes of intrinsically generated coupling: synchronization of neural phase, the correlation of amplitude envelopes and the coupling between phase and amplitude ^9–13^. In addition, these coupling modes operate in multiple frequency bands and scales, mediating different cognitive functions ^6,7,9,14–17^, and providing different channels for long-range communication ^18,19^. Phase coupling, in particular, has been suggested to serve the routing of information through cortical networks and to promote selective communication between distant brain areas ^6,7,20^, as well as to mediate stimulus selection and prediction ^6,7,17,19,21^. However, how phase coupling supports the brain communication across scales, and how it relates to behavior, is not well understood.

A natural scenario to test the role and mechanisms of phase coupling in brain dynamics at multiple scales are multisensory tasks. In the human brain, studies of multi-scale network dynamics have remained challenging although substantial advances have been made using non-invasive methods such as EEG or MEG. In particular, only a few studies have addressed the relation between phase coupling and multisensory processing ^22,23^. In animal studies, the potential role of functional connectivity involved in multisensory processing has so far been investigated only by multielectrode recordings from the same cortical areas ^24,25^ or by simultaneous recordings from at most two different regions ^26^. Short-range neural interactions are well established at the level of microcircuits ^27–29^. Invasive studies on functional connectivity between distant population in behaving animals typically have recorded simultaneously only from relatively small numbers of sites ^20,30–33^.

Electrocorticographic (ECoG) arrays offer an option to study simultaneously the relation between multiple-scales of network dynamics and behavior ^18,34–36^. Because of their bio-compatibility and recording stability, they have been used for the study of functional connectivity in networks underlying cognitive and sensorimotor functions ^37–41^ in humans ^11,42^, monkeys ^18,38^ and rodents ^43^. Furthermore, small-sized ECoG electrodes can provide local information at the level of cortical columns ^34,44^ which, when distributed across distant regions in an array, provide a unique opportunity to study the interaction between local and global cortical dynamics.

Here, we investigated how changes in functional connectivity and power, indicative of global and local dynamics, respectively, determine the detection of multisensory stimuli. We hypothesized that in a multisensory detection task the functional coupling between modalities, rather than local power, determines the performance. We tested this hypothesis in a large-scale cortical network involving visual, auditory, somatosensory and parietal areas using a custom-made 64-channel ECoG array in behaving ferrets. Animals were trained in a 2-alternative-forced-choice paradigm to detect brief auditory and visual stimuli presented either left or right from the midline. A key finding of our study is that phase coupling of visual and auditory regions with parietal cortex was predictive of task performance. Our results suggest that fluctuations in the networks state, particular with respect to long-range connectivity, are decisive for the response of the upcoming task during multisensory processing.

## Materials and Methods

### Animals

All experiments were approved by the independent Hamburg state authority for animal welfare (BUG-Hamburg) and were conducted in accordance with the guidelines of the German Animal Protection Law. Data were recorded in four adult female ferrets (Mustela putorius furo; Euroferret, Dybbølsgade, Denmark) aged 2 and 4 years (n=2 each). Animals were individually housed in standard ferret cages equipped with an enriched environment, under controlled conditions (21°C, 12-hour light/dark cycle, with lights on at 8:00 a.m.). The animals had ad libitum access to food pellets, while access to tap water was restricted for 8 hours prior to the experiments and training procedures. All behavioral testing was conducted during the light cycle, specifically between 10:00 a.m. and 2:00 p.m. The animals were treated with a deslorelin acetate (4.7 mg implant) chip to control their ovarian activity^45^. Animals were trained on a two-alternative forced-choice task, described in detail below, involving lateralized detection of visual and auditory stimuli that we have established previously ^46^. After completion of all recording sessions (for details, see below), animals were euthanized in deep anesthesia with a lethal dose of pentobarbital (400 mg/kg i.p.). Subsequently, animals were transcardially perfused with fixative (4% paraformaldehyde in phosphate-buffered saline) and brains were removed for histological processing.

### Sensory stimuli

All experiments were carried out in a dark sound-attenuated chamber (Acoustair, Moerkapelle, Netherlands). Visual and auditory stimuli were generated with the Psychophysics Toolbox ^47^. Visual stimuli were presented on an LCD monitor (Samsung SyncMaster 2233, frame rate 100 Hz) placed 20 cm in front of the animal. Auditory stimuli were generated digitally with a sample rate of 96 kHz and delivered through two Beyerdynamic T1 speakers. Auditory stimuli consisted of white noise bursts of 100 ms duration, including 5ms cosine rise/fall, presented with intensities between 2 and 62 dB SPL. Visual stimuli consisted of circular gratings (22.5 °, 0.2 cycles / °, 5 Hz) with Michelson contrast (Cm) between 0 and 0.38. A static random noise pattern located in the center of the screen was presented to indicate trial onset by a decrease in contrast (not overall luminance). Multisensory stimuli consisted of both stimulation types presented simultaneously.

### Training

Initially, ferrets were handled and accustomed to a housemade flat-bottomed tube, which served as the animals’ enclosure during the experiment. Figure 1A shows a schematic of the experimental setup. Subsequently, animals were trained in the spatial detection task, during which the unimodal as well as crossmodal psychophysical thresholds were estimated. In the task, we used a crossmodal approach that bears on established unimodal training paradigms ^48^. Animals were restricted from access to water for a period of 8 hours before the measurements, and conditioned by using water rewards during the task. In the first phase of the study, the animals were trained to detect unimodal auditory and visual stimuli presented in a randomized fashion. Auditory and visual unimodal detection thresholds were determined using 20 different stimulus amplitudes (auditory: 2-62 dB SPL; visual: 0-0.38 Cm) in a 1 down / 3 up staircase procedure ^49^. Next, unimodal and bimodal thresholds were assessed in a combined approach, using the previously determined unimodal thresholds to adjust the test parameters (for details of the procedure see ^46^). Subsequently, the ferrets were accustomed to electrophysiological recordings during the detection task with a reduced set of stimulus amplitudes (eight per modality), adjusted to the individual psychometric functions to acquire a higher number of trials in the performance range of interest around 75 % accuracy.

**Figure 1.**
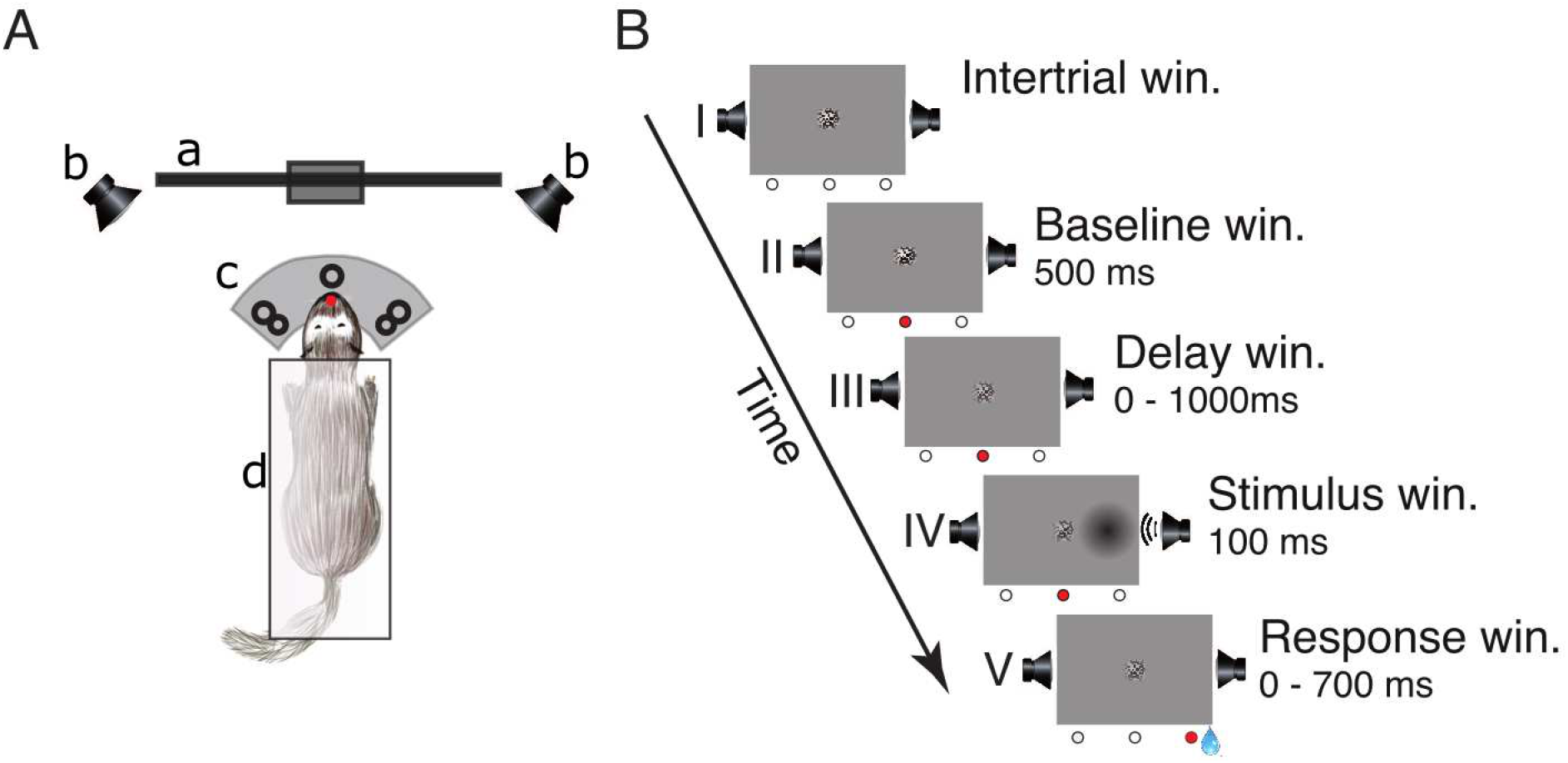
Experimental setup and lateralized detection task. (A) Schematic of the experimental setup in a top view: a) LCD-screen, b) speakers, c) three light-barrier-waterspout combinations (left, center, right; the red dot indicates a broken light-beam), d) pedestal. (B) Sequence of events in a single trial: (I) inter-trial window, (II) baseline window, (III) delay window, (IV) stimulus window, and (V) response window. The three circles below each frame represent the state of the light-barriers (white = unbroken, red = broken). The center of the screen displays a static circular random noise pattern.

### Detection task

The trial schedule of the detection task is shown in Figure 1B. To initialize a trial, the ferret had to maintain a central head position to break the central light-barrier for 500 ms. This caused the static random noise pattern in the center of the screen to decrease in contrast informing the animal that the window for stimulus presentation (from 0 to 1000 ms after trial onset) had started. During this interval, the animal had to further maintain its central head position. A stimulus was presented for 100 ms on either the left or the right side. Stimulus presentation could be unimodal visual (‘V’), unimodal auditory (‘A’) or a temporally congruent bimodal combination of an auditory and a visual stimulus presented on the same side. This combination either consisted of a variable auditory stimulus accompanied by a visual stimulus of constant contrast (‘Av’) or, conversely, a visual stimulus of varying contrast supported by an auditory stimulus of constant amplitude (‘Va’). The intensity value of the accompanying second stimulus with constant amplitude was set at a level of 75% accuracy. After stimulus offset, the animal had to respond within 700 ms by moving its head to the stimulated side; otherwise, the trial was considered as a miss (no response). If the response was correct the animal received a water reward (∼80 μl) from a spout at the stimulus position and could immediately start the next trial. If the response occurred prematurely (before stimulus onset or within 100 ms after stimulus onset), was incorrect (wrong side) or omitted, the trial was immediately terminated, followed by a 2000 ms interval during which no new trial start could be initialized.

### ECoG array implantation

Micromachining technology ^35^ was used to design and implement an ECoG array that matched the anatomy of the ferret brain (Fig. 2A and B). The thin-film (10 µm) polymide-foil ECoG contained 64 platinum electrodes with a diameter of 250 µm each, arranged in a hexagonal formation at an inter-electrode distance of 1.5 mm (Fig. 2C and 2D).

**Figure 2.**
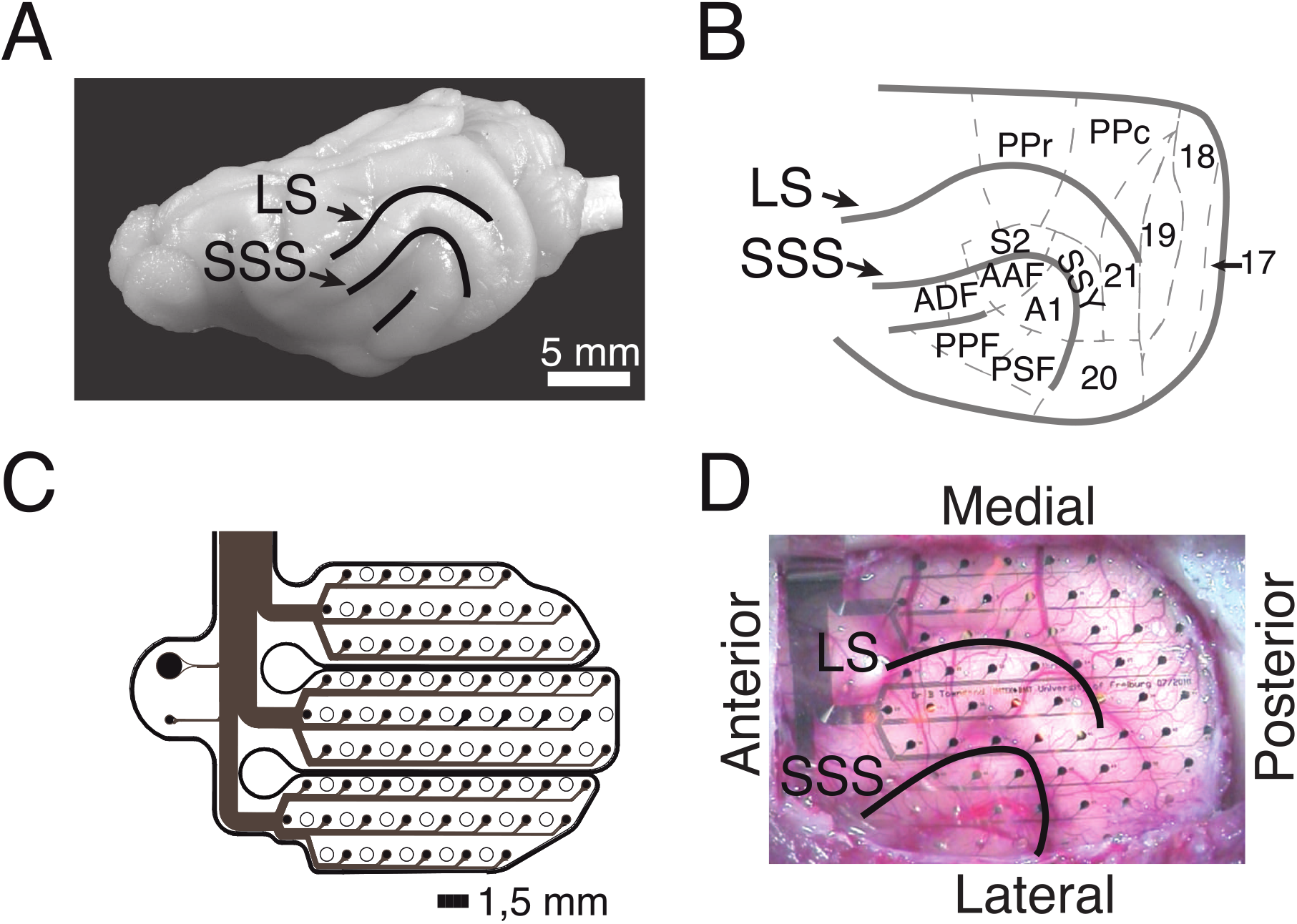
ECoG recordings from the ferret brain. (A) Lateral view of the left hemisphere. Lines depict sulci in the posterior part of the hemisphere. (B) Anatomical organization of the posterior part of the ferret brain (adopted from Bizley et al., 2007). (C) Schematic of the implanted ECoG (contact spacing: 1.5mm; ø: 250µm). (D) In-situ picture of the ECoG array over the left posterior hemisphere during implantation. Abbreviations: LS, lateral sulcus; SSS, suprasylvian sulcus; S2, secondary somatosensory cortex; PPr/c, rostral/caudal posterior parietal cortex; 17, 18, 19, 20, 21, early and higher-order visual cortex; SSY, suprasylvian field; A1, primary auditory cortex; AAF, anterior auditory field; ADF, anterior dorsal field; PPF, posterior pseudosylvian field; PSF, posterior suprasylvian field.

The surgical procedure for the implantation of the ECoG array started with the induction of general anesthesia by using a ketamine-medetomidine mixture (10 mg/kg and 1 mg/kg, respectively). During surgery, anesthesia was maintained by ventilating the animal with isoflurane (1-1.5%) in combination with 70% N_2_0 and 30% O_2_. To monitor the state of anesthesia, physiological parameters such as the electrocardiogram (ECG) and rectal temperature were monitored throughout the surgery. All surgical procedures were performed under sterile conditions. After the operating field was prepared, a craniotomy was performed using a saline-cooled ultrasonic microsaw (Mectron) in order to expose the posterior half of the left cerebral hemisphere. The dura was carefully removed and the ECoG array was gently placed on the surface of the cortex such that it covered occipital, temporal and parietal areas (Fig. 2D). The dura was then folded back over the ECoG array and an absorbable artificial dura was placed above the ECoG, covering the trepanation to full extent. The excised piece of bone was fixed back in place with titanium plates and screws and subsequently the gaps were filled with fast set absorbable bone putty. Finally, the ECoG’s connector was placed on top of the skull and fixed in place with titanium screws and dental acrylic. After the surgery, the animals received analgesic (carprofen, 2.5 mg/kg) and antibiotic (enrofloxacin, 5 mg/kg) medication. To align electrode positions across animals, the ECoG array placement and the cortical parcellation introduced by Bizley et al. (2007) was utilized (Fig. 2B). For each animal, the exact ECoG array position over the posterior cortex was photographically documented during surgery. The locations of all 64 ECoG electrodes were then projected onto a scaled illustration of a ferret model brain. Each electrode was then mapped to the underlying cortical area.

### Electrophysiological recording and preprocessing

Local field potentials (LFPs) were digitized at 1.4 kHz and sampled simultaneously with a 64- channel AlphaLab SnR recording system (Alpha Omega Engineering, Israel). The high pass filter was set at 0.1 Hz and the low pass filter at 357 Hz.

To ensure comparability across animals and modalities, only recording blocks with an accuracy of 75±10% were considered in all electrophysiological analyses. Furthermore, this selection of trials ensured better comparability between all four modalities because the value that was fixed during the bimodal stimulation was set at a level of 75% accuracy for all stimulation amplitudes. To assess a sufficient amount of trials with stimulus presentation contralateral to the implanted hemisphere per modality in the 75% accuracy range, trials collected on different days were pooled; see ^46^ for non-stationarity effects across sessions.

All offline data analysis was performed using custom scripts in Matlab (The Mathworks Inc, MA). The hardware-filtered LFP signals were band-pass filtered with a phase-preserving fourth-order Butterworth filter between 2 and 200 Hz. Next, band-stop Butterworth filters at 49-51, 99-101 and 149- 151 Hz were applied to notch the line noise. Subsequently, the continuous LFP signals were cut into epochs aligned to trial, stimulus and response onset, respectively. In each of these analysis time windows data were cut from 500 ms pre- to 500 ms post-onset, with 500 ms real data padding on each side to prevent edge artifacts in frequency space. Afterwards, we applied independent component analysis (ICA)^50^ to the concatenated analysis window data in order to detect and remove ECG, muscle and eye blink artifacts. Next, we re-referenced the LFP time series of each ECoG contact with the LFP time series of its next posterior neighbor. This processing step created 55 virtual electrodes from the 64 recorded contacts. The new virtual electrode was located in the midpoint between both real electrode positions. Subsequently, the oscillatory signal components of unimodal and bimodal trials where analyzed using spectral decomposition.

### Spectral power analysis

Channel-wise spectral power was computed by taking the square of the absolute value of the Fourier transform. Spectra were computed for all three analysis time windows using a Hanning window approach (2-200 Hz, 2 Hz steps, 500 ms window; for the LFP spectrogram the window was shifted from – 500 ms to 500 ms around the window of interest) in 1 ms steps.

### Quantification of phase locking

To estimate phase synchronization between ECoG signals, we calculated the phase locking value (PLV)^51^ across all frequencies (2-200 Hz). In general, the instantaneous phase θ was extracted from the analytic signals that were produced by the Fourier transform of the convolution of the ECoG time series with the Hanning window. The PLV between channels A and B at carrier frequency *f* is defined as follows:

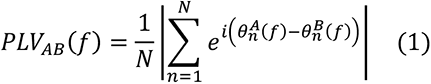

For an event related paradigm the circular average of phases is in general non-vanishing, and a non-vanishing result using the classical definition of PLV does not imply statistical dependence between phases in two electrodes. Therefore, we corrected for evoked effects and defined the PLV here as

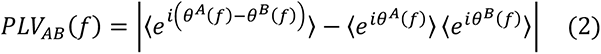

Where 〈 〉 denotes average over trials. For an infinite number of trials, the PLV defined here vanishes exactly if the phase is electrode A and B are statistically independent. To contrast PLV between modalities, we first computed the global mean PLV for the same time windows selected in the spectral power analysis. Subsequently, the PLV between anatomical regions, as defined in Figure 2B, was calculated by averaging across all virtual electrodes overlaying the same area. To normalize and compare PLV values from hit and miss trials we computed the sensitivity index (d’), which is defined by:

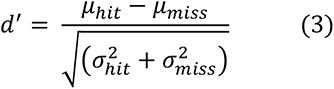

where *μ* denotes the mean and *σ* the standard deviation for hit and miss trials, respectively.

### Principal component analysis of functional connectivity

We used a similar approach to that of Leonardi et al. ^52^ to identify hidden sub-networks during the baseline period that may be associated with performance by applying principal component analysis (PCA) to trial -based connectivity matrices. To facilitate the contrast in the analysis we separated matrices in groups of hits and misses. Before applying PCA, the global mean connectivity matrix was subtracted from matrices of all trials within each group. To avoid redundancy we took only the lower triangular part of each *n*x*n* connectivity matrix (excluding elements of the diagonal) and then vectorized it to obtain a vector of *n(n-1)/2* elements. The vectors of all animals were concatenated along the trial dimension into an array of *(n-1)/2* x*T* elements, with *T* being the total number of trials. PCA was then performed on the resultant matrix. Eigenvectors or principal components (PCs) can be understood as features or subnetworks that characterize the variation across the collection of correlation matrices and represent network patterns. Because these capture connectivity patterns they are also called eigenconnectivities ^52^. Finally, to reduce dimensionality and obtain the main eigenconnectivities, matrices were thresholded by the lowest and strongest 10% with respect to their mean.

### Directionality analysis

To estimate the direction of causal influences we calculated the phase slope index PSI (Ψ) ^53^. PSI is highly robust against false estimates caused by confounding factors of very general nature, and is defined as:

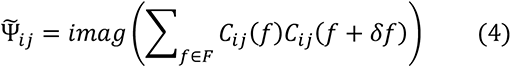

where 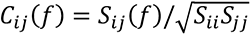 is the complex coherency, *S* the cross-spectral matrix, δ*f* the frequency resolution. *F* is the set of frequencies over which the slope is summed. The complex coherency ***C*** is a Hermitian matrix and consequently PSI is antisymmetric and, hence, directional. Intuiti vely, PSI is the slope (as a function of frequency) of the phase difference between two signals averaged within a frequency band. The slope is larger, when the time lag between the signal and the coupling increases. The sign depends on which signal is earlier. It is constructed such that it vanishes (apart from stastical fluctuations) if the observed signal is an instantaneous superposition of independent sources, and the average across frequencies is weighted with coupling strength. To normalize 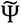 we divided it by its standard deviation

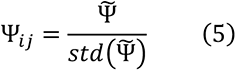

stimated by the jackknife method. The acquired PSI matrices where thresholded equivalently to the eigenconnectivity matrices and their sign was extracted.

## Results

To investigate neural responses and task-related fluctuations of functional connectivity, we recorded local field potentials (LFPs) via 64-channel ECoG arrays chronically implanted in four adult female ferrets.

### Multisensory effects on behavior

Before the electrophysiological recordings were conducted, a psychometric evaluation was performed on each animal to determine the stimulus parameters (see Methods). Figures 3A and B show examples of psychometric functions fitted to the behavioral data for the *A* and *Av*, and for the *V* and *Va* conditions, respectively. For further analysis, data were taken only from trials with stimulus intensities at threshold to allow comparisons across conditions and animals (Fig. 3A, B).

**Figure 3.**
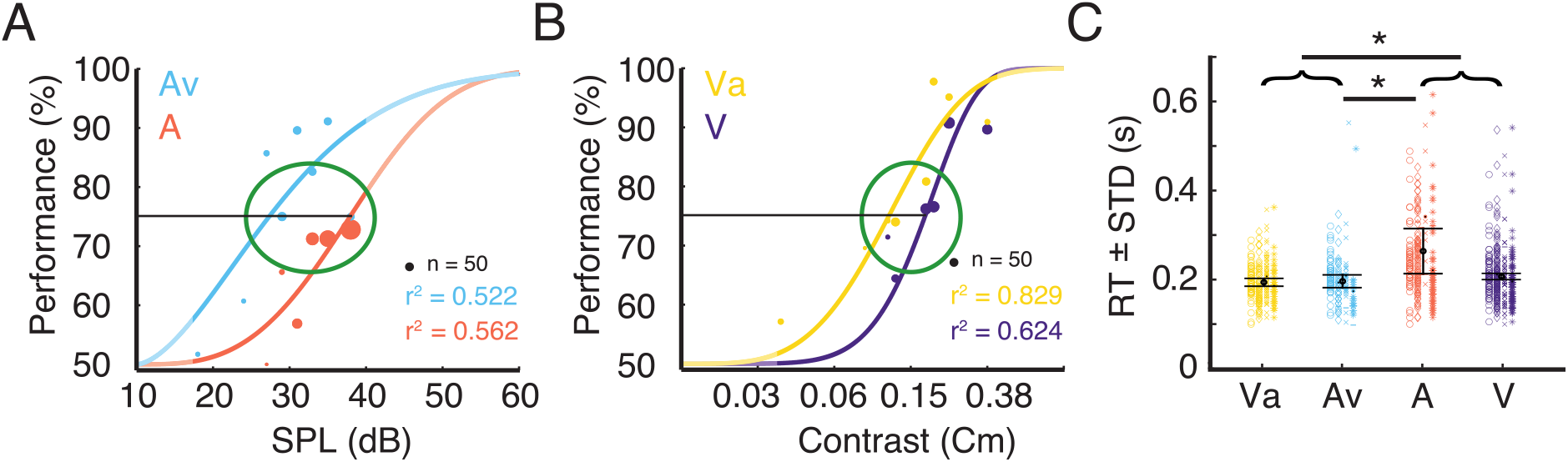
Trial selection for electrophysiological analysis and corresponding RTs. (A) Example data for task performance from one ferret. The plot shows psychometric functions for the uni- and bimodal condition. Stimulus response function fitted to performance data for the eight different stimulus intensities for the unimodal auditory condition (A) and the condition where the variable auditory stimulus was accompanied by a visual stimulus of constant contrast (Av). The dot diameter indicates the number of trials at a given stimulus intensity. The green ellipse indicates the stimulus intensity range around 75% performance from which trials were selected for subsequent analysis. The unmasked parts of the graphs indicate the range of the actually tested stimulus amplitudes. (B) Data from the same animal as shown in A for the unimodal visual condition (V) and the condition where the variable visual stimulus was accompanied by an auditory stimulus of constant intensity (Va). (C) Reaction times (mean ± standard deviation) for the four different conditions for the selected trials. The four different shapes (circle, diamond, cross, star) represent the single trial RT of each animal. Error bars indicates the mean RT±STD across animals. Black asterisks indicate significant differences between modalities or pooled data from bimodal (Va and Av) and unimodal (A and V) stimulus pairs (brackets).

Reaction times (RT) have been broadly used to quantify differences between unimodal and multisensory stimulation. A *t*-test was run to investigate the differences between bi- and unimodal conditions. Responses to bimodal stimulation were, on average, faster than those to unimodal stimulation (*p* < 0.05). To evaluate whether this effect was driven by the auditory or the visual condition, we computed a one-way analysis of variance (ANOVA) on the mean RT of each animal and condition, with sensory modality as a main factor. This revealed a main effect of condition (*F* (3, 12) = 5.3, *p* < 0.05), but post hoc *t*-tests only revealed significant differences between the *A*-trials and both bimodal conditions. The population averages of RTs for *Av*, *Va* and *V* were not significantly different (Fig. 3C). Nevertheless, the population mean RT of *V* (0.207 sec) was slightly higher than both, *Va* (0.194 sec) and *Av* (0.197 sec). The significant difference between the RTs in the *A* and the *Av* condition suggest a multisensory interaction effect leading to reduction in RT for the bimodal stimulation. However, our data do not suggest a substantial improvement of visual detection by a concurrent auditory stimulus.

### LFP power reflects stimulus processing and response preparation

A major aim of our analysis of the electrophysiological data was to test the relation of power and functional connectivity to task performance by contrasting hit and miss trials. In the hit trial group, we included all trials with stimulus intensities around 75% threshold and stimulation contralateral to the implanted ECoG (265 ± 14 hit trials per animal). To match the number of hit trials we used all miss trials (236 ± 45 miss trials per animal) throughout all sessions (15 ± 2 sessions per animal). Miss trials were defined as those with sensory stimulation that did not elicited a behavioral response, i.e., the ferret maintained the centered head position throughout the response window. The number of false responses, i.e., orientation of the ferret to the side contralateral to the sensory stimulation, were too few and highly variable across animals (104 ± 58 false responses per animal) and, thus, were not considered for analysis of the electrophysiological data.

We checked whether the spectral characteristics of LFPs immediately before task execution predicted the animal’s performance. To assess how LFP spectral characteristics evolve during hit, miss and bi- and unimodal stimulation, we computed the power across time windows aligned to baseline, stimulus presentation and response onset for each animal individually (Fig. 4). Results of all animals were pooled after correction for different trial numbers. Power differences between hit and miss trials were analyzed in the theta (4-8 Hz), alpha (8-16 Hz), beta (18-24 Hz), gamma (30-80 Hz) and high-gamma (80 - 160 Hz) frequency band across the analysis time windows (Tab. 1). In addition, one-way ANOVAs were calculated on power values within each frequency band with condition as the main factor to examine differences related to stimulus type (Fig. S1; Suppl. Tab. 1).

**Figure 4.**
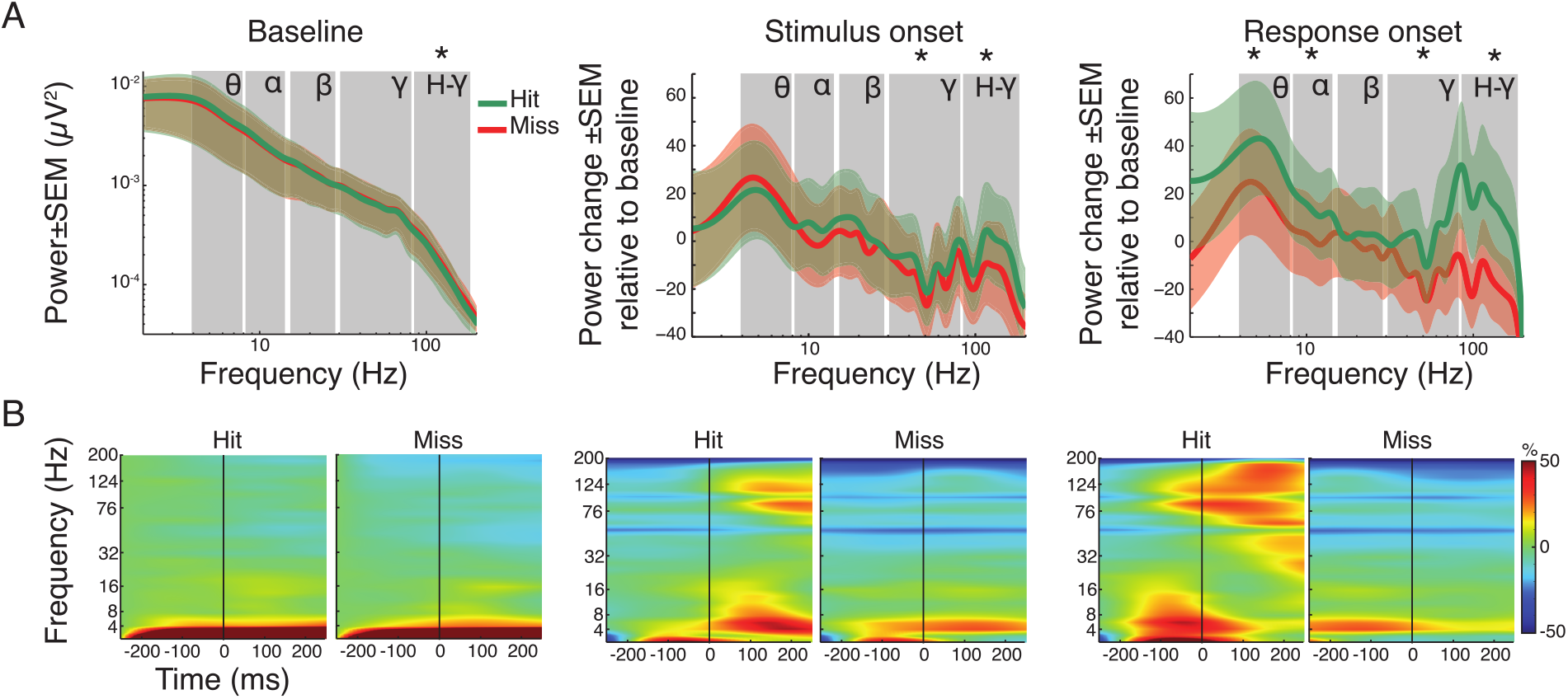
Spectral power analysis of pooled hit and miss trials. (A) Grand average power spectra for all hit (green) and miss (red) trials±standard error of the mean (SEM). For the baseline window absolute power is shown. Note that for the time windows around stimulus and response onset, the spectral changes shown are relative to the baseline window. Asterisks indicate significant differences between hits and misses within the specified frequency band (FDR corrected). Labels indicate theta, alpha, beta, gamma and high-gamma band, respectively. (B) Time-frequency representation of power in the three analysis time windows, expressed as change relative to baseline before trial onset for all hit and miss trials. The vertical line represents trial, stimulus and response onset, respectively.

Figure 4A illustrates the grand average spectra, while Figure 4B shows the time-frequency representation of power changes during each time window (baseline, stimulus and response onset) for both hit and miss trials. During the baseline, only the high-gamma frequency-band (80 - 160 Hz) showed moderated but significant differences, with higher power for miss trials (*p* < 0.05; FDR corrected). Comparison of the average spectra in the stimulus onset analysis window showed significant reductions for hit and miss trials in the gamma frequency-range (30 - 160 Hz) compared to baseline (Tab. 1). However, hit trials showed significantly higher power compared to miss trials after stimulus onset in both gamma (*p* < 0.001) and high-gamma (*p* < 0.001) frequency bands (Fig. 4A). Around response onset, hit trials consistently showed significantly higher power compared to miss trials in the theta- (*p* < 0.01), alpha- (*p* < 0.01), gamma- (*p* < 0.001) and high-gamma band (*p* < 0.001). Statistical comparison across time windows within hit- and miss-trials revealed significant decrease and increase in the gamma frequency range compared to baseline and stimulus onset, respectively (Tab. 1; all *p*-values, within and across windows of interest, FDR corrected).

Supplementary Figure S2 shows the topographies of the respective spectral differences between hit and miss trials. In the baseline window, power topographies showed high spatial uniformity across all frequency bands. Around stimulus onset, notable differences occurred mainly in the beta band where, in hit trials, higher beta power differences occurred in parietal areas. The highest regional differences occurred in the response window. Occipital regions exhibited higher power across all frequency bands during response onset in miss trials compared to hit trials. In contrast, auditory and parietal areas showed increased power in lower frequency bands (theta and alpha) for hit trials. An increase of gamma band power occurred in the response window for hit trials in regions around the lateral sulcus (Fig. S2).

To investigate stimulus-type dependent effects within hit-trials a one-way ANOVA between the four conditions (*Av*, *Va*, *A* and *V*; Fig. S1; Suppl. Tab. 1) was calculated. It showed no main effect in the factor condition across frequency bands during baseline, stimulus or response onset. This suggests that power changes related to stimulus processing and response preparation did not depend on the modality of the presented stimulus or on crossmodal interactions.

### Functional connectivity predicts performance

To determine whether functional connectivity predicts the animals’ performance we computed the PLV between all pairs of ECoG channels for the different frequency bands. PLV analysis was performed for the same time windows as the power analysis: aligned to baseline, stimulus and response onset. Figure 5 displays the population average PLV spectra for all hit and miss trials as well as the difference between hits and misses (Fig. 5B) and the relative spectral changes in the stimulus and response onset window relative to baseline (Fig. 5B). Paired *t*-tests were applied between hit and miss trials within and across all time windows for each frequency band separately. Within each analysis window, significantly higher PLV was observed for hits compared to miss trials in all frequency bands, with exception of the theta band during response onset (*p* < 0.05, FDR corrected; Fig. 5A).

**Figure 5.**
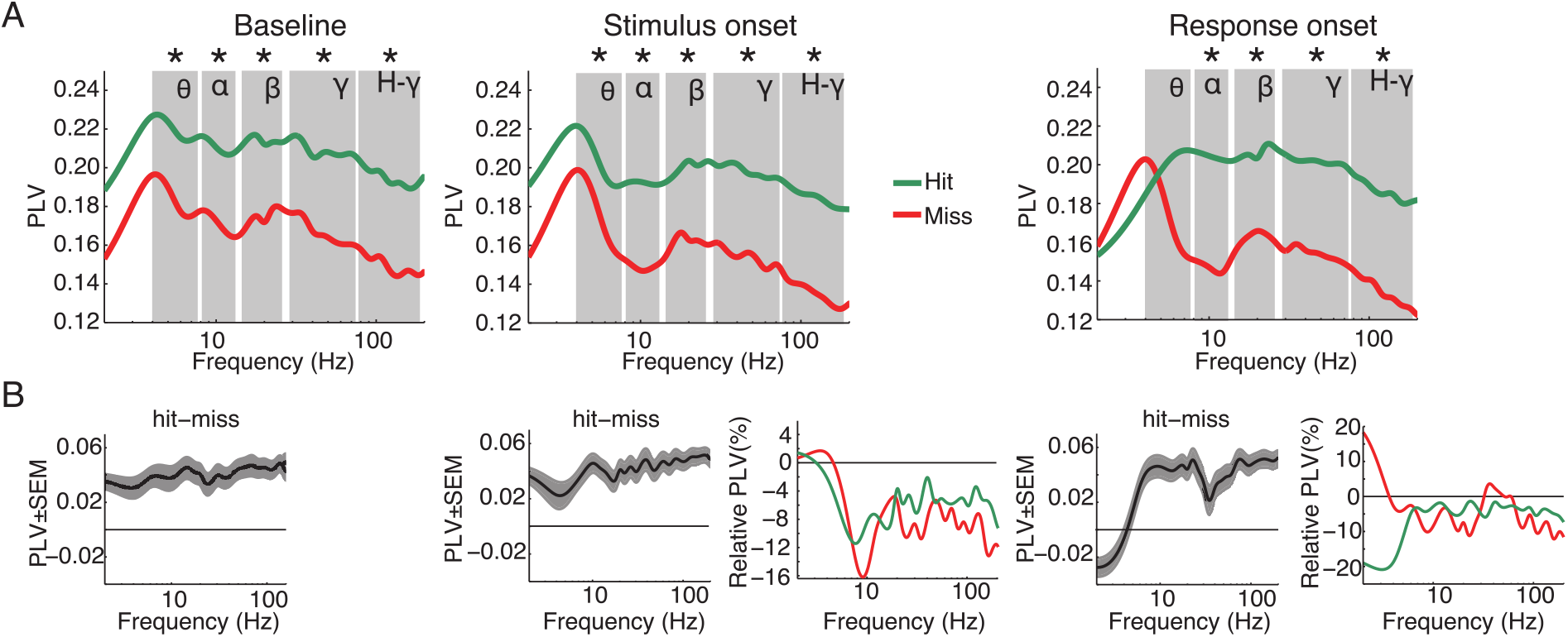
Grand average of functional connectivity for hit and miss trials. (A) Grand average phase locking value (PLV) for all hit (green) and miss (red) trials for the analysis time windows (baseline, stimulus and response onset). (B) Differences between hits and misses for the three analysis windows and changes of PLV around stimulus and response onset, expressed as changes relative to the baseline window. Asterisks indicate significant differences between hits and misses within the specified frequency band (FDR corrected). Labels in A denote theta, alpha, beta, gamma and high-gamma band, respectively.

When compared to the baseline time window, PLV was significantly reduced in the stimulus and response onset windows in nearly all frequency bands. Exceptions were in the theta frequency band for miss trials around stimulus as well as response onset.

A one-way ANOVA within analysis time windows and frequency bands with stimulus condition as the main factor revealed significant differences in PLV between stimulation conditions (*Av*, *Va*, *A* and *V*; Fig. S3). During the baseline period there was no significant effects in the main factor condition in the theta frequency band (*F* (3, 44) = 2.46, *p* > 0.05). However, starting with the alpha (*F* (3, 76) = 4.98, *p* = 0.003), beta (*F* (3, 60) = 3.71, *p* = 0.016), gamma (*F* (3, 412) = 31.89, *p* < 0.001) and high-gamma (*F* (3, 652) = 56.99, *p* < 0.001) frequency bands, there was a main effect of conditions. This effect persisted also for the windows around stimulus onset (theta: *F* (3, 44) = 1.23, *p* > 0.05; alpha: *F* (3, 76) = 3.74, *p* = 0.015; beta: *F* (3, 60) = 3.5, *p* = 0.021; gamma: *F* (3, 412) = 26.12, *p* < 0.001; high-gamma: *F* (3, 652) = 55.67, *p* < 0.001) and response onset (theta: *F* (3, 44) = 2.31, *p* > 0.05; alpha: *F* (3, 76) = 3.63, *p* = 0.017; beta: *F* (3, 60) = 3.67, *p* = 0.017; gamma: *F* (3, 412) = 25.12, *p* < 0.001; high-gamma: *F* (3, 652) = 51.24, *p* < 0.001). Furthermore, post-hoc *t*-test confirmed a constant pattern of significant differences between conditions (Suppl. Tab. 2). In particular, for hit trials connectivity was higher in the *A* and *Av* conditions than in the *V* condition, and there was a trend for connectivity in bimodal *Va* trials to be higher than in unimodal *V* trials (Fig. S3).

Taken together, in contrast to the power, the PLV revealed significant differences between hit and miss trials. These differences occurred over a broad range of frequencies across all analysis time windows. These results suggest that large-scale phase-fluctuations in network state are determinant to the animals’ performance.

### Large-scale coupling shows specific changes during the task

To examine functional connectivity in relation to the topography of cortical areas, we assigned the data of each ECoG contact to a distinct region based on the functional map of cortical areas from Bizley et al.^54^ (see Methods) and constructed functional connectivity matrices accordingly (Fig. 6). Cortical areas were grouped in three functional systems, comprising visual (areas 17, 18, 19, 20, 21), auditory (areas A1, AAF, ADF, PPF and PSF), and parietal areas (SSY, PPc, PPr, S2). Differences between hit and miss trials were expressed using the sensitivity index (d’).

**Figure 6.**
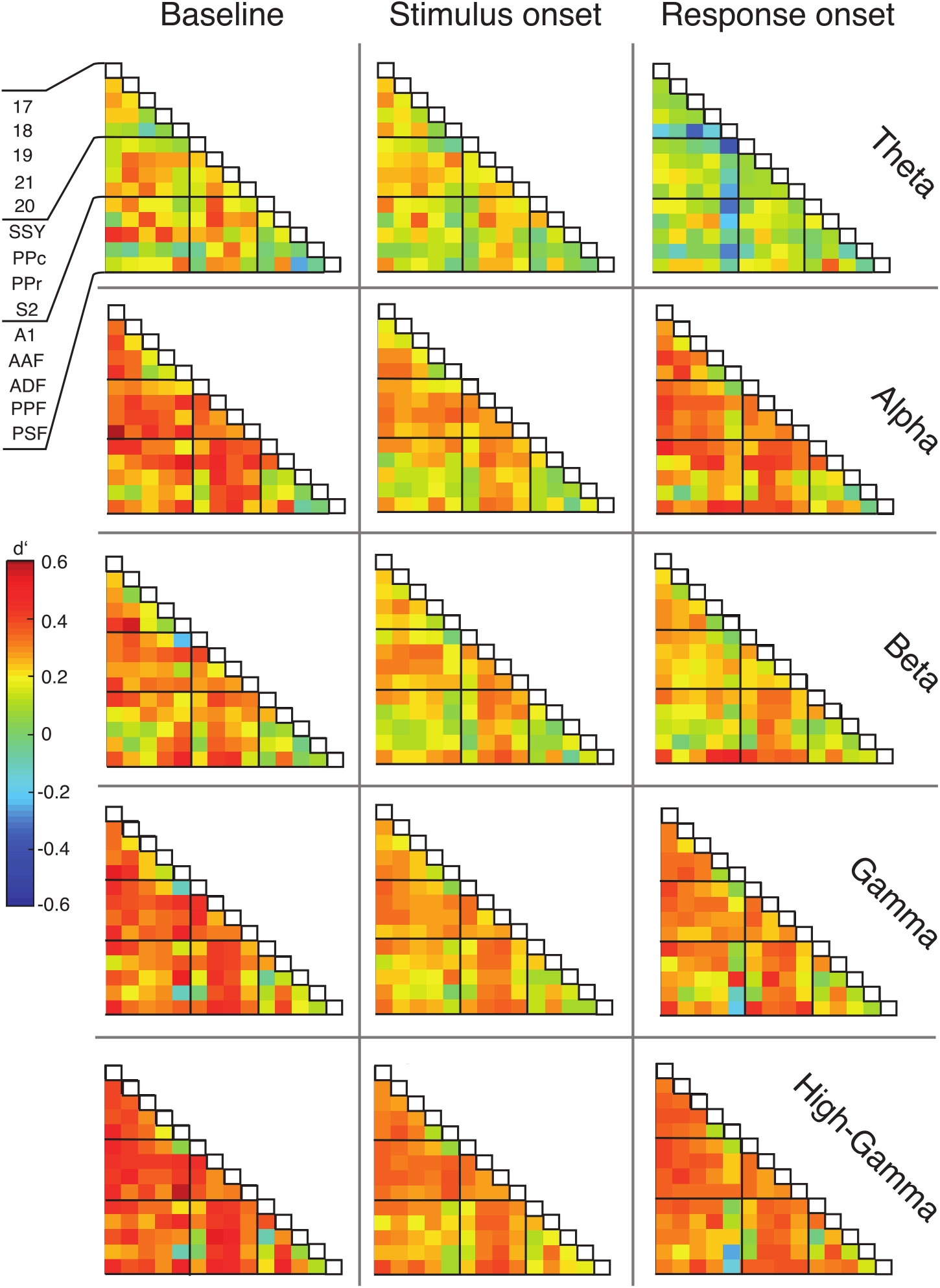
**Matrices of functional connectivity across anatomical areas**. Cells show population average phase locking values (PLV) for the different frequency bands, expressed as d’ for hits vs. misses, for all analysis time windows. Abbreviations as in Fig. 2.

In line with the grand average PLV results, the connectivity matrices generally displayed positive d’ values and, thus, higher functional connectivity for hit compared to miss trials. (Fig. 6). This effect was most prominent during the baseline window and decreased during the stimulus presentation. For the alpha, gamma and high-gamma band, the PLV difference between hit and miss trials increased again in the response window. Generally, connections between early sensory cortices and parietal cortex displayed the highest differences between hit and miss trials, in particular during the baseline window (Fig. 6; Fig. S5).

### Eigenconnectivity reveals subnetworks related to performance

To detect the subnetworks that most strongly contributed to differences in the animal’s performance, we examined the baseline period on a single-trial basis by performing PCA for hits and misses along the trial dimension (see Methods). We focused on a subset of areas (18, 19, 21, PPc, PPr, A1 and AAF) from the recorded functional systems for the analysis of task-related subnetworks, to reduce dimensionality and to examine which functional connections were most strongly related to hits versus misses. The sign of the interactions was extracted from PSI matrices to derive directionality of the coupling in subnetworks.

Figure 7 shows directed eigenconnectivity patterns in the beta and gamma bands for hit and miss trials in the first and second PCs. Only those connections which had the smallest and the largest differences relative to the mean connectivity in the baseline window are displayed (complete matrices and explained variance for PCs 1-6 and all frequency bands are shown in Fig. S4). PC1 did not show substantial differences between hits and misses (Fig. 7A; Fig. S4). For both hits and misses, the first PC revealed stronger interactions between auditory and visual areas, but weaker interactions between visual and parietal cortices. In contrast, PC2 revealed substantial differences in functional connectivity between hit and miss trials. These differences were most prominent in the theta, alpha and beta band, but occurred also in the gamma band (Fig. 7B; Fig. S4). In the beta band, coupling within the visual system was stronger for hit compared to miss trials, whereas coupling of auditory area AAF with parietal and visual areas was stronger for hits than for misses (Fig. 7B). Interestingly, an increase in connectivity between AAF and visual areas was a common pattern in miss trials for both PC1 and PC2 in the beta and gamma band (Fig. 7). The third to sixth PCs revealed more complex interaction patterns, with increases as well as decreases for functional connections in both hit and miss trials (Fig. S4).

**Figure 7.**
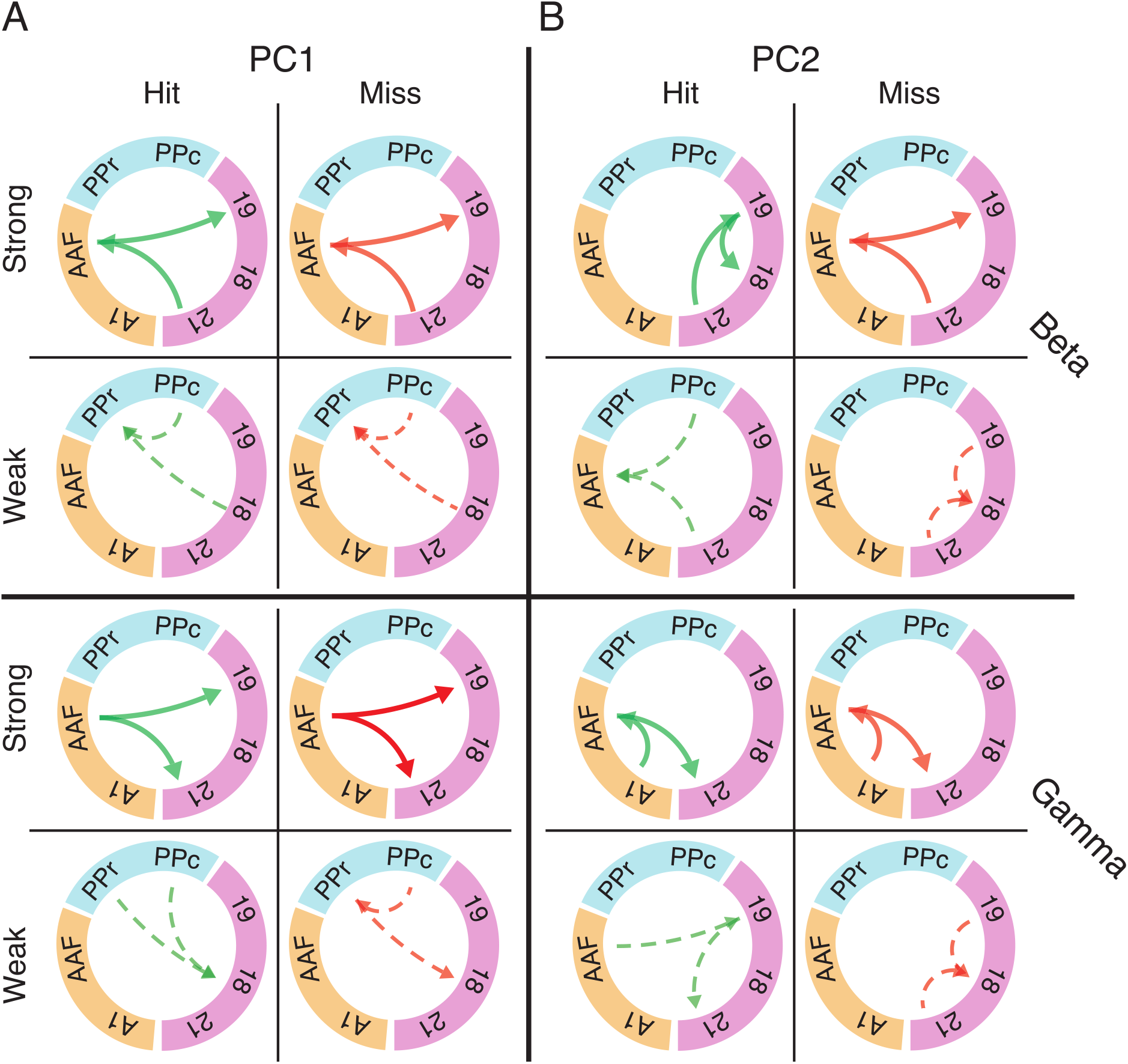
Eigenconnectivity (EC) of hit and miss trials for the baseline window. Depicted are EC networks for beta and gamma of the first two PCs, which together explain 42% and 32% of the variance in the PCA for hits and misses, respectively. Green and red arrows represent hit and miss networks, respectively. Thick solid lines indicate the upper 10%, and thin dashed lines the lowest 10% of connections in the EC distribution. Arrowheads indicate the direction of information flow extracted by PSI. Colored circle segments represent the three functional systems (visual = magenta; parietal = blue; auditory = orange). Note that EC matrices for the first 6 PCs with all connection are shown in Fig. S4. Abbreviations as in Fig. 2.

It should be noted that the eigenconnectivity patterns cannot be directly compared to the global connectivity differences between hits and misses described in the preceding sections, because they reflect changes in subnetworks, relative to the average network coupling strength, in the two groups of trials. More importantly, our analysis shows that changes in connectivity do not occur arbitrarly across the network, but they are distributed in subnetworks that allow a coexisting increase/decrease in phase coupling.

### Between- and within-system interactions differ in their relation to behavior

The topography of functional connectivity changes was further analyzed by contrasting between-system and within-system interactions (Fig. 8; Fig. S5). A previous study from our group showed that this is a suitable classification to determine global/local effects associated to changes in brain state (Stitt et al. 2017). For hit trials, interactions between the three functional systems were, on average, stronger than functional connections within the respective system. Regarding the within-system connectivity, the auditory system showed the lowest d’. In the period from baseline to stimulus onset, the strength of d’ decreased across all frequency bands. With response onset the spectral profile of d’ became more heterogeneous. In the theta-band d’ decreased further, whereas it remained unaltered in the beta-band, and increased in the alpha-, gamma- and high-gamma band.

**Figure 8.**
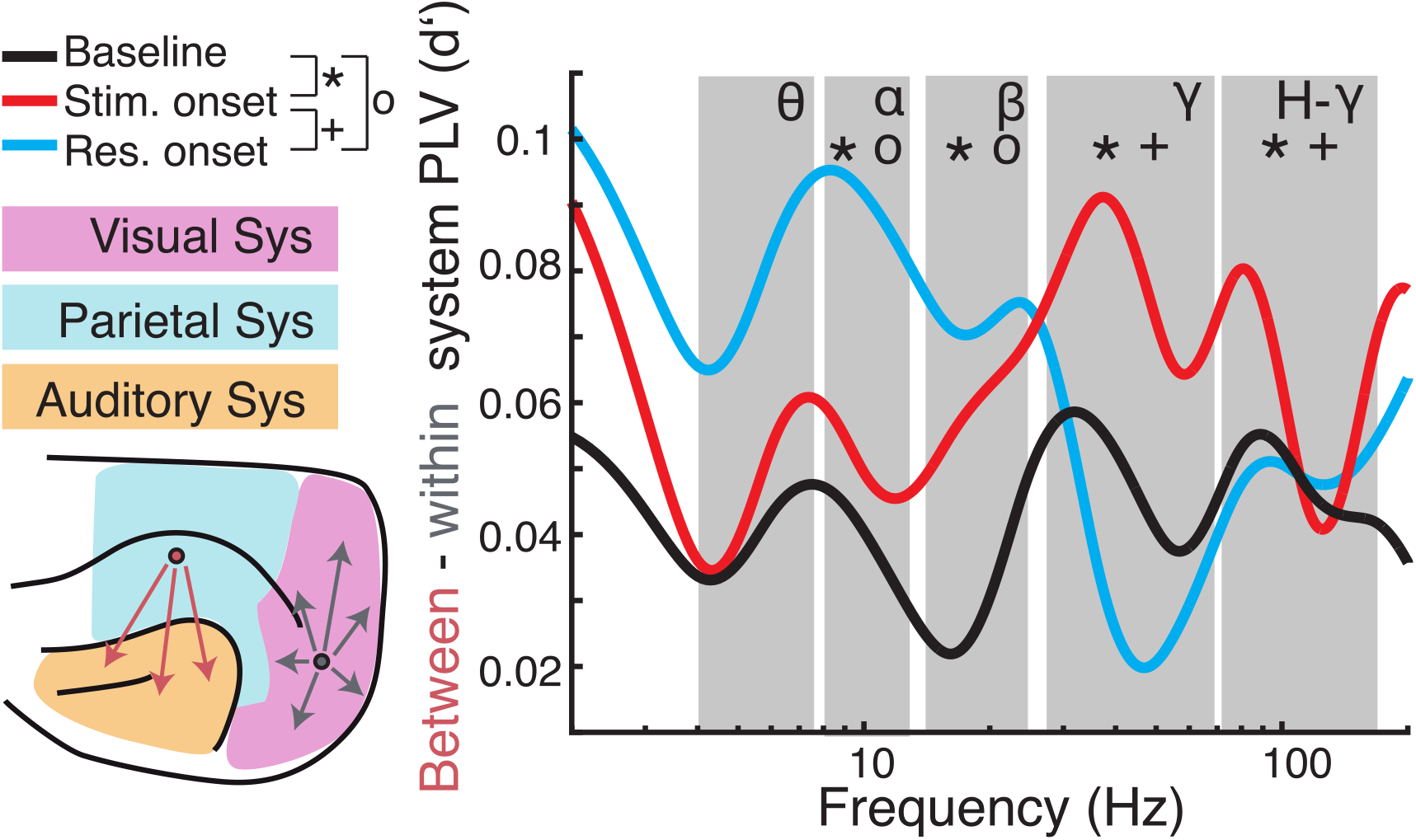
Comparison of long-range (between-systems) and short-range (within-system) functional connectivity. (Left) Cortical regions were grouped into visual (magenta), parietal (blue), or auditory (orange) systems. PLV was averaged for all pairs of regions that were located within the same system (grey arrows) and pairs between systems (red arrows). Subsequently, d’ was calculated for hit and miss trial PLVs. (Right) Differences of between-region and within-region PLV (expressed as d’) are plotted as a function of carrier frequencies. Large values indicate higher between-system PLV during hit compare to miss trials. Symbols indicate significant differences between baseline and stimulus onset (*), baseline and response onset (o), and stimulus onset to response onset (+), respectively.

For statistical analysis, we averaged PLV d’ values and derived the difference of d’ values between long-range (between-system) and short-range (within-system) connectivity and calculated statistics across analysis time windows for each frequency band (Fig. 8; for statistics across systems and within analysis time windows and frequency see Fig. S5; for condition specific statistics see Fig. S6). To reveal dynamic effects within frequency bands of interest a one-way ANOVA with time window as the main factor was computed. There was no main effect in the theta band (*F* (2, 6) = 4.39, *p* > 0.05). The ANOVA exposed significant effects for the main factor time window (alpha (*F* (2, 12) = 17.17, *p* < 0.001; beta (*F* (2, 9) = 14.43, *p* = 0.002; gamma (*F* (2, 75) = 3.71, *p* < 0.001; high-gamma (*F* (2, 120) = 17.94, *p* < 0.001). Post hoc *t*-tests revealed significant differences between baseline and stimulus onset for all frequency bands with the main effect (alpha: *p* = 0.011; beta: *p* = 0.002; gamma: *p* < 0.001; high-gamma: *p* < 0.001), between stimulus and response onset for the gamma frequency range (gamma: *p* = 0.009; high-gamma: *p* < 0.001) and between baseline and response onset in the alpha- (*p* < 0.001) and beta- (*p* = 0.006) frequency bands (Bonferroni corrected).

Overall, comparison of functional connectivity between and within cortical systems across the different analysis time windows revealed that between- compared to within-system connectivity was stronger during the task period compared to the pre-stimulus baseline. In hit-trials, phase coupling was higher between systems in the gamma frequency range during stimulus onset compare to baseline and response onset. Furthermore, low frequency (alpha and beta) coupling between systems increased around response onset in hit trials compared to miss trials. These results suggest that fluctuations in the network state, particularly with respect to long-range connectivity, are related to task performance.

## Discussion

This study has aimed at investigating the cortical network dynamics at multiple spatial scales and subnetworks. We hypothesized that in tasks that involve multiple distant modalities the functional connectivity, rather than the local activity, becomes a main determinant for performance. We addressed this question in an audiovisual detection task in ferrets. We used an ECoG approach to record a large-scale network comprising visual, auditory, somatosensory and parietal areas in behaving animals over extended time periods.

In line with our hypothesis, our data show that pre-trial baseline connectivity predicts the animals’ performance, suggesting that fluctuations in coupling across the network lead to variability in behavior. In contrast, mean power in the pre-trial baseline was not associated to performance. Analysis of the topography of connectivity differences between hits and misses revealed specific patterns, reflecting different functional subnetworks, associated to the response. In particular, higher global phase coupling of visual and auditory regions to parietal cortex was predictive of task performance, yet a reduction in connectivity in certain subnetworks may be necessary for distinct functional systems to detect stimuli appropriately. We also observed that long-range coupling became more predominant during the task period compared to the pre-stimulus baseline and changed its spectral profile over the course of the trials.

### Relating large-scale connectivity to task performance

Among the established functional connectivity measures, coherence, in particular, has been associated to cognitive tasks in humans involving perceptual selection ^55^, attentional selection ^19,56^, working memory ^57^ or speech processing ^23,58^. Animal studies have demonstrated a relation of long-range phase coupling in various frequency ranges to sensorimotor processing, attentional selection and working memory ^18,30,32,33,38,59–61^.

We found that phase coupling, in contrast to local power, differed already between hit and miss trials in the pre-trial baseline. The data indicate that stronger phase coupling in the network may lead to higher efficiency of stimulus processing and response preparation. This is in line with earlier studies that have demonstrated that increased synchrony enhances communication across brain areas (e.g., in the context of attention ^7,38^) supporting the role of functional connectivity in linking task-relevant modalities to areas involved in selection of responses ^9,31^. Indeed, we find that stronger phase coupling of early sensory to parietal areas was predictive of task performance.

Contrary to earlier studies, our results appear not to correspond to the typical vigilance- or attention-related signatures in pre-stimulus functional connectivity characterized by increases in high frequency and decreases in low frequency bands ^38,56,62^. Although the average connectivity differences between hits and misses had a rather broad spectral profile, analysis of the ratio between long-range (between cortical systems) and local interactions (within cortical systems) revealed spectral shifts in this ratio between the task period and the pre-stimulus baseline. This result highlights the role of within-area high-frequency coupling during stimulus processing, which may reflect stronger bottom-up signaling. This could relate to attentional gating, allowing relevant information to pass through the network to higher areas. In contrast, the predominance of low-frequency interactions in the epoch around response onset might indicate stronger top-down information flow ^18^. Alternatively, in that interval of the trial, parts of the network could already return to a default state dominated by low frequency oscillations. Averaged across areas, spectral differences between connections with dominant bottom-up and top-down information flows, might level out and, thus, yield the broad spectral profile of connectivity changes observed in the present analysis.

### Functional connectivity and multisensory processing

As discussed above, dynamic functional coupling likely constitutes a mechanism for integration of distributed neural signals and sensory systems ^5–7,63,64^. It has been suggested that similar mechanisms might operate for the integration of information across different sensory systems. Thus, multisensory interactions might involve dynamic coupling of oscillatory signals arising in different cortical systems ^8,65^. In the human brain, the vast majority of studies on neural oscillations and crossmodal processing have focused on local power changes, and only few investigations have addressed the relation between multisensory processing and functional coupling ^22,23^. Using a data-driven approach for analysis of functional coupling in the human EEG, we could recently demonstrate that coherence in networks involving parietal and sensorimotor areas predicts performance in a visuotactile matching task ^66^.

Simultaneous recordings from auditory cortex and superior temporal sulcus in the monkey have revealed increased coherence during congruent auditory and visual stimulation ^26,67^. While these in-vivo studies investigated multisensory interactions only by recordings from the same cortical areas or by simultaneous recordings from at most two different regions, our study has addressed multisensory networks at a larger scale involving visual, auditory and parietal cortical regions. While our data clearly demonstrate a relation of functional connectivity to task performance, we did not obtain evidence for stimulus-specific changes of coupling between visual and auditory areas, resembling recent results in anesthetized ferrets ^13^, where modification of coupling structure occurred only on longer time scales and not in a stimulus-related manner.

The absence of profound connectivity effects related to multisensory interaction may relate to the nature of the task used in our study, which did not require integration of features across modalities, but only the rapid detection of highly transient stimuli. Nonetheless, our results on connectivity data support the notion that significant functional coupling can occur already between early sensory areas, suggesting that multisensory integration can already occur at early processing stages and does not solely rely on binding of information at higher processing levels ^8,24,54^. This is in line with anatomical data suggesting that direct projections from primary auditory to visual cortices can enable functional coupling supporting early multisensory interactions ^68^.

### Network state and behavioral variability

A key result of our study is that phase coupling differed between hit and miss trials already in ongoing activity in the baseline period. The local power in this period did not differ between hits and misses. Our data suggest that fluctuations in the network state occur which lead to variability in the animals’ behavior, and that these state changes are primarily reflected in shifts of long-range connectivity, rather than changes in the dynamics of local populations. Early studies assumed that ongoing neural activity corresponds to noise resulting from random signal fluctuations without any functional relevance ^69,70^. However, this view has been challenged by evidence showing that ongoing activity carries information that can shape sensory and cognitive processing ^6,13,22,71–73^. Such fluctuations of ongoing activity do not occur only locally, but are strongly synchronized across spatially distributed neuronal populations ^13,36,74,75^.

Phase coupling of oscillations in pre-stimulus epochs has been shown both in animal and human studies to predict perception and performance in cognitive tasks. For instance, studies in monkey visual cortex indicate that fluctuations in gamma-band phase coupling modulates the speed at which animals can detect a behaviorally-relevant stimulus change ^76^. EEG studies in humans provide convergent evidence that pre-stimulus fluctuations in phase coupling can modulate target detection ^62^. Furthermore, intrinsic fluctuations of phase coupling are associated with fluctuations in perceptual states in ambiguous stimulus settings. Fluctuations in beta-band or gamma-band phase coupling have been shown to predict the perceptual state in ambiguous sensory paradigms ^55,77^ and decision making in near-threshold stimulation regimes ^78^. Our current results corroborate and extend this evidence by showing that, at perceptual threshold, detection of the lateralized stimuli by the animals is biased by phase coupling in the pre-stimulus interval.

A crucial element in our approach is the ability to simultaneously monitor numerous cortical areas, enabling us to quantify within-system and between-system interactions and to characterize these in a spectrally resolved manner. Our analysis showed that, in particular, long-range connectivity between different functional systems was related to successful stimulus detection. This is in line with the hypothesis that long-range phase coupling may serve efficient transmission of task-relevant information in sensorimotor networks ^6,7,79^. One important finding was that the variations in connectivity patterns associated to changes in behavioral performances, do not occur arbitrarily along the whole cortex, but in specific subnetworks, which were identified by extracting principal components of functional connectivity. Whether these subnetworks are associated to a decision making network, or rather sensory processing is not clear. However, since most of the nodes include primary sensory areas, and the connectivity changes occurred before the stimulus was presented, we speculate that the first reason for miss trials is a sub-optimal function of stimulus coding.

Our results also support the view that the characterization of changes in brain-state strongly benefits from inclusion of connectivity analyses. The observation that changes in network state are reflected, in particular, in fluctuations of large-scale connectivity, is in line with results of other recent studies. In rats^80^ has being shown that functional connectivity within and between brain areas is modulated across behavioral states in a region-specific manner. Supportive evidence has been obtained in studies on the human brain ^81,82^. Using the same ECoG recording approach for the study of ongoing activity, we have observed that functional connectivity shows state-dependent reconfiguration which can also involve shifts in the ratio between short- and long-range interactions ^36,75^. These findings raise the question of possible mechanisms that might modulate large-scale functional connectivity in a state- and task-dependent manner. Possible candidates are changes in the output of ascending neuromodulatory systems ^83^. Phase coupling has long been known to be influenced by neuromodulators^84^. For instance, activation of cholinergic brain stem nuclei has been shown to enhance gamma-band phase coupling in cortical networks ^85^. Neuropharmacological evidence suggests, furthermore, that noradrenergic brain stem inputs can modulate large-scale functional connectivity in cortex ^86^.

## Conclusion

Our study has demonstrated functional coupling across visual, auditory and parietal areas during a lateralized detection task in the ferret. Analysis of power for hit and miss trials revealed significant differences around stimulus and response onset. In contrast, phase coupling already differed between hits and misses at baseline, suggesting an essential role of large-scale functional connectivity in the animal’s ability to perform the task. In particular, higher phase coupling of visual and auditory regions to parietal cortex was predictive of task performance. We observed that long-range coupling became more predominant during the task period compared to the pre-stimulus baseline. Taken together, these results suggest that fluctuations in the network state, particular with respect to long-range connectivity, are critical determinants of the animals’ behavior. Future studies might address the relation to the underlying structural connectivity and the mechanisms that give rise to the observed variability of phase coupling.

## Acknowledgements

We would like to thank Dorrit Bystron for her assistance in the experiments. This research was supported by funding from the DFG (SFB936-178316478-A2, SPP1665-EN533/13-1, SPP2041-EN533/15-1, A.K.E.; SFB936-178316478-Z3, G.N.) and by the European Union (project cICMs, ERC-2022-AdG-101097402, A.K.E.). Views and opinions expressed in this paper are those of the authors only and do not necessarily reflect those of the European Union or the European Research Council. Neither the European Union nor the granting authority can be held responsible for them.

## Data Availability

Results data obtained for this manuscript can be provided upon request from the corresponding authors.

## Additional Information

They are no competing interests.

## Author Contributions

K.J.H, F.P. and A.K.E. conceived and designed experiments. G.E. wrote animal ethics permission. K.J.H and F.P. surgically implanted arrays. K.J.H. performed experiments. K.J.H., F.P., E.G.L. and G.N. provided analyzing tools. K.J.H. analyzed data. K.J.H., E.G.L. and A.K.E. interpreted data and wrote the paper. A.K.E. and G.N. acquired funding. All authors edited and approved the manuscript.

