## Supplementary material for "Dynamic changes in large-scale functional connectivity prior to stimulation determine performance in a multisensory task": 3 supplemental tables and 6 supplemental figures.

### Dynamic changes in multiscale functional connectivity prior to stimulus determines performance in a multisensory task

### (Supplement)

#### Tables

**Table 1. Statistics for frequency resolved power analysis for hit and miss trials.** The table shows *p*-values for all *t*-tests within frequency bands, trial outcome, and across all time points of interest; *italics* indicate significant comparisons ($p<0.001$; FDR corrected).

| Frequency | Theta | Alpha | Beta | Gamma | High-Gamma |
|---|---|---|---|---|---|
| | Baseline vs. stimulus onset window | | | | |
| Hit vs. hit | 0.22 | 0.26 | 0.44 | *>0.001* | 0.053 |
| Miss vs. miss | 0.23 | 0.54 | 0.99 | *>0.001* | *>0.001* |
| | Stimulus onset vs. response onset window | | | | |
| Hit vs. hit | 0.03 | 0.18 | 0.08 | *>0.001* | *>0.001* |
| Miss vs. miss | 0.21 | 0.62 | 0.53 | 0.08 | *>0.001* |
| | Response onset window vs. baseline | | | | |
| Hit vs. hit | 0.10 | 0.46 | 0.98 | 0.01 | *>0.001* |
| Miss vs. miss | 0.25 | 0.58 | 0.79 | *>0.001* | *>0.001* |

**Table 2. Statistics for frequency resolved functional connectivity analysis for all time points of interest for hit and miss trials.** The table shows *p*-values for all *t*-tests within frequency bands, trial outcome and across all time points of interest; *italics* indicate significant contrasts (FDR corrected).

| Frequency | Theta | Alpha | Beta | Gamma | High-Gamma |
|---|---|---|---|---|---|
| | Baseline vs. stimulus onset window | | | | |
| Hit vs. hit | 0.12 | *<0.001* | *<0.001* | *<0.001* | *<0.001* |
| Miss vs. miss | 0.75 | *<0.001* | *<0.001* | *<0.001* | *<0.001* |
| | Stimulus onset vs. response onset window | | | | |
| Hit vs. hit | 0.03 | *<0.001* | 0.36 | *<0.001* | *<0.001* |
| Miss vs. miss | 0.79 | 0.07 | 0.67 | 0.17 | *<0.001* |
| | Response onset window vs. baseline | | | | |
| Hit vs. hit | *<0.001* | 0.02 | 0.003 | *<0.001* | *<0.001* |
| Miss vs. miss | 0.84 | *<0.001* | *<0.001* | *<0.001* | *<0.001* |

#### Figures

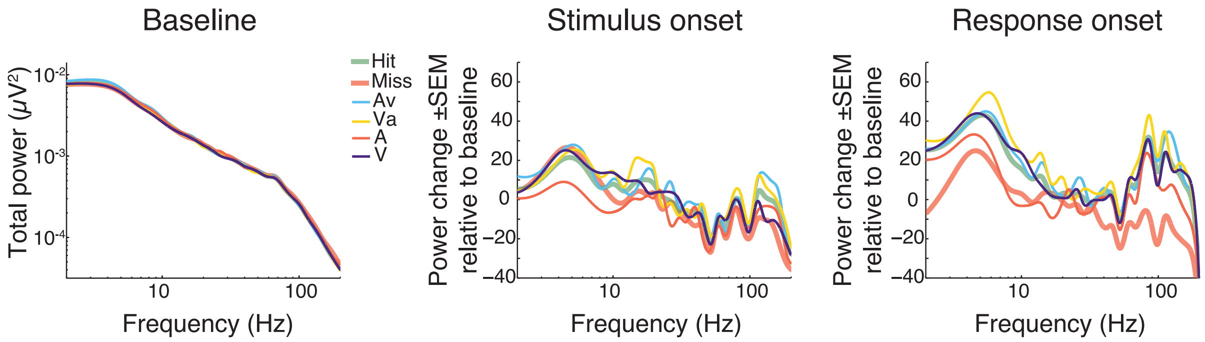


Figure S1. Left: total power in baseline intervals for each condition and response. Middle and right plots show the power change relative to baseline and response onset, respectively, for each condition.

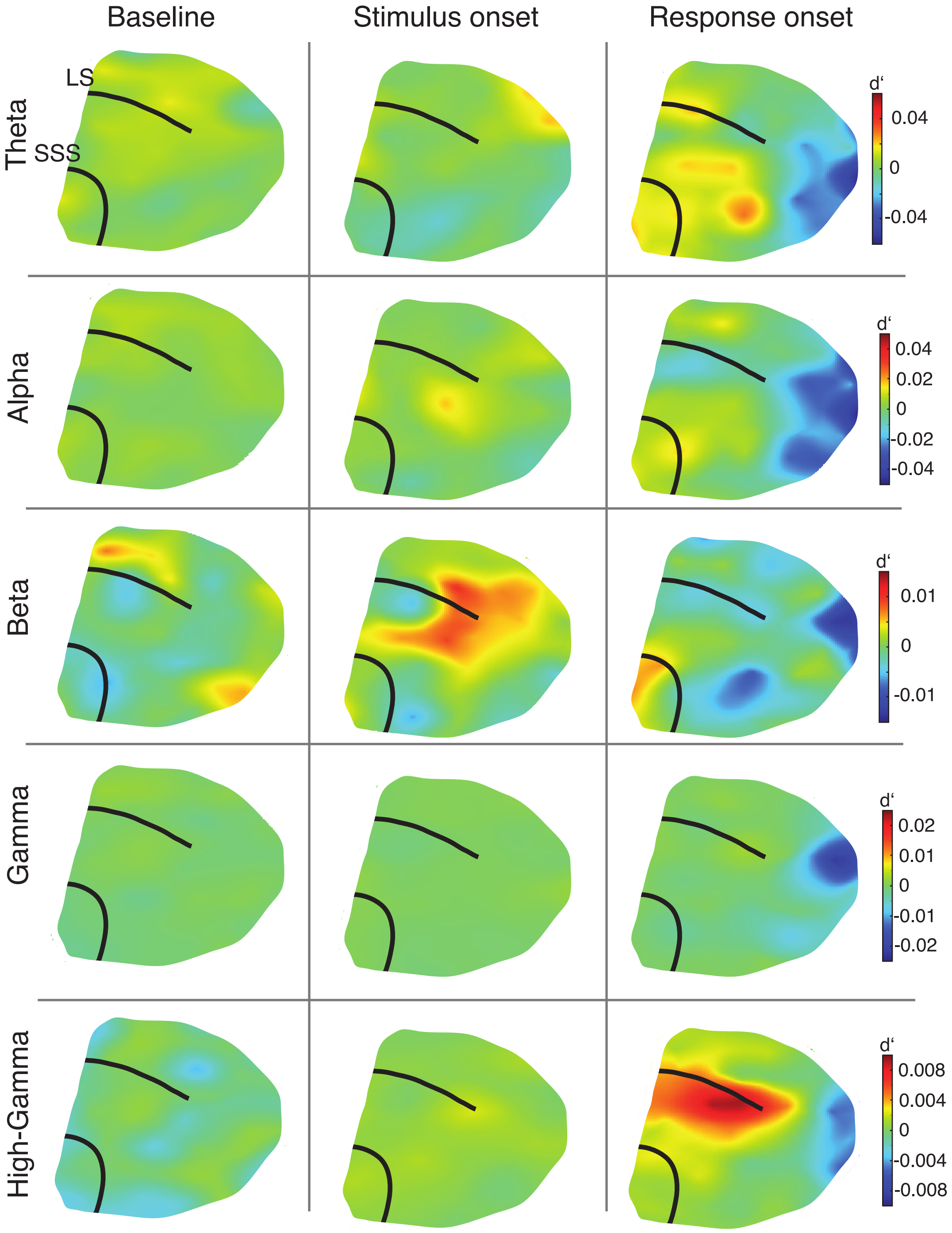


Figure S2. Topographic distribution of spectral differences between hit and miss trials for three different time windows (columns) and frequency bands (rows).

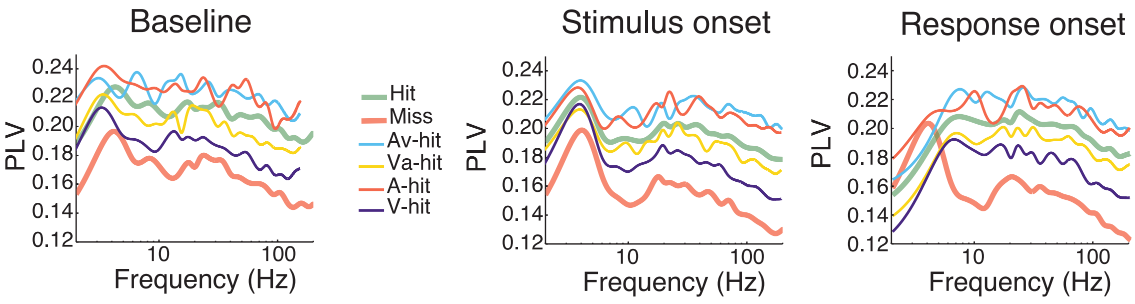


Figure S3. Effects of stimulus conditions and response on phase connectivity (phase-locking value) at three different time windows. In contrast to power (Fig S1), differences in connectivity are observed in the baseline period.

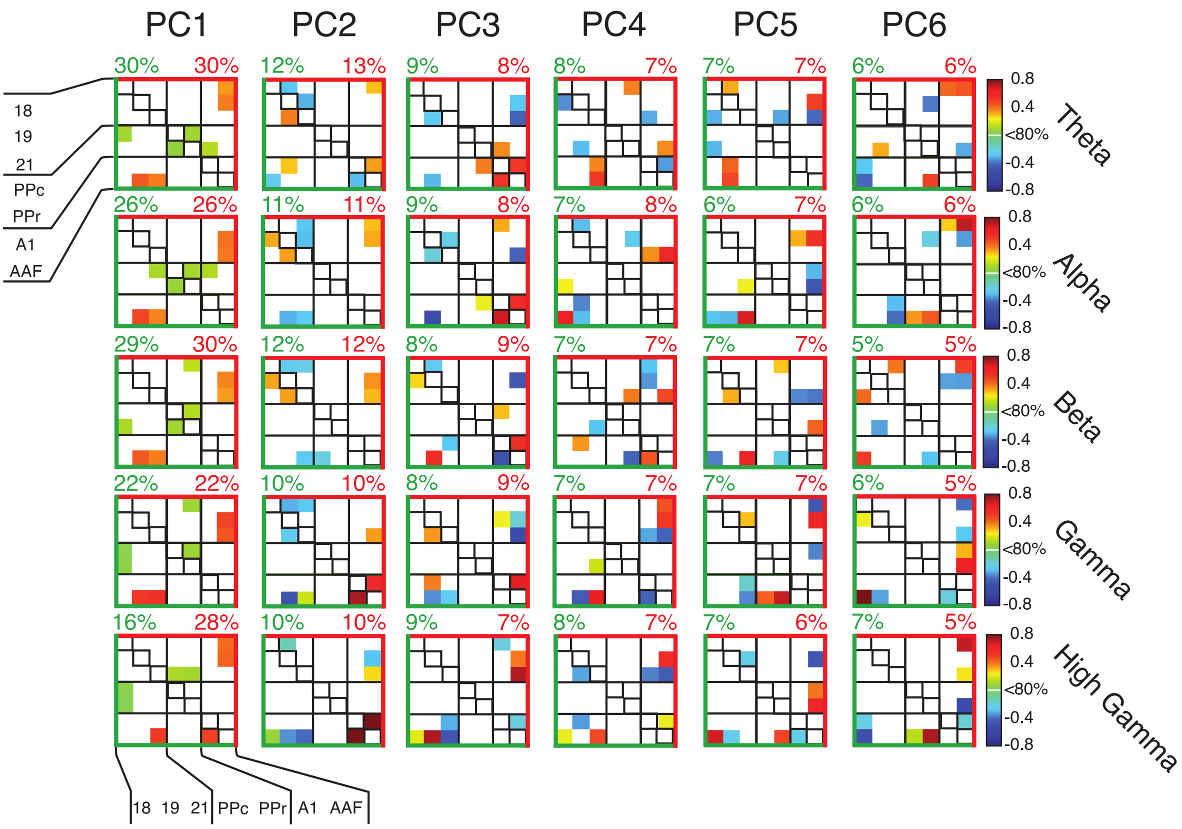


Figure S4. Eigenconnectivity patterns for hit trials (low triangle – green) and miss trials (upper triangle – red). Displayed are the connections that became stronger (hot colors) or weaker (cold colors) relative to the means connectivity.

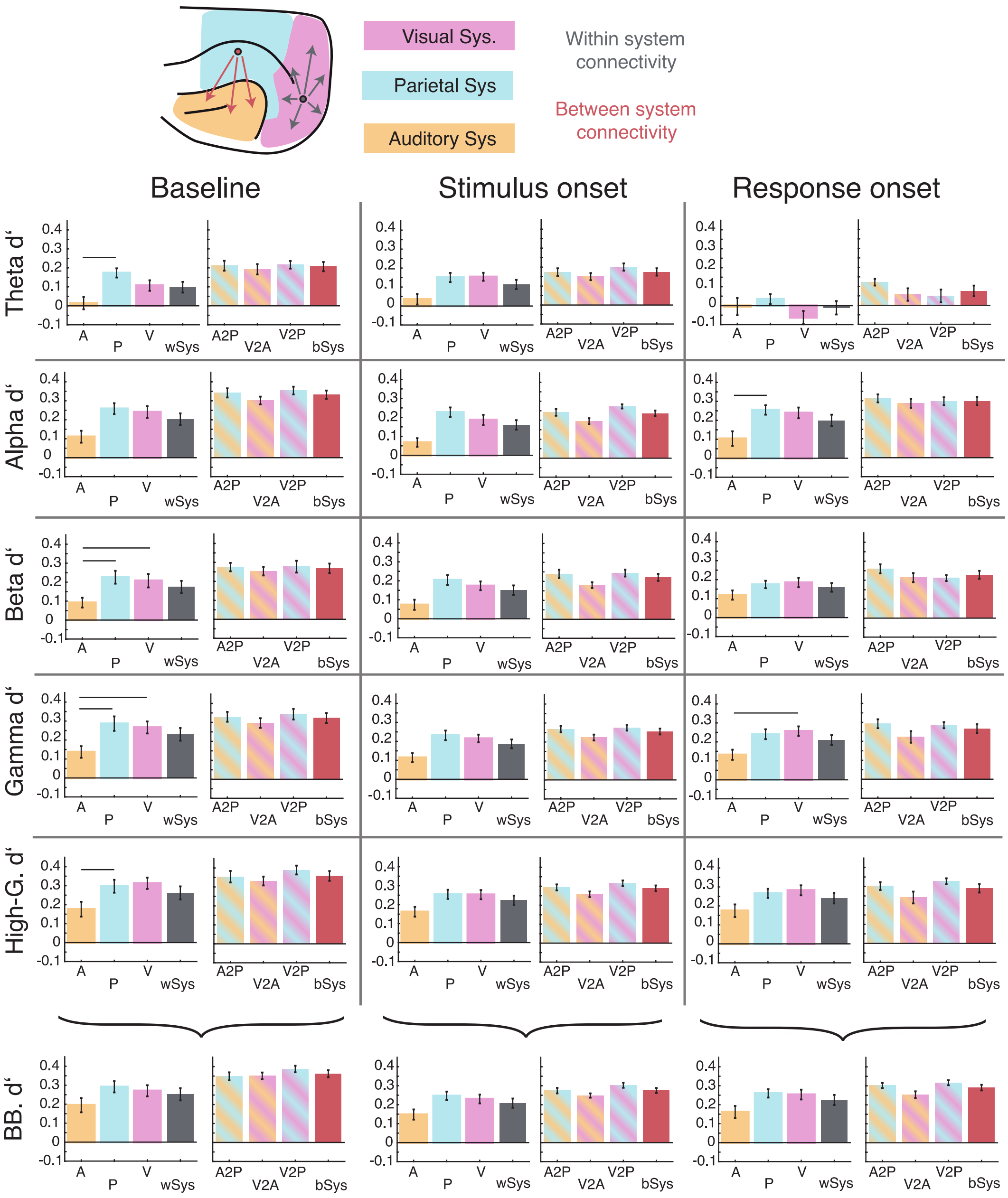


Figure S5. Contrast of the interactions in within-system (full colors) and between-system (patterned colors). Bars in the between-system condition are represented by the colors of the individual system. Horizontal lines mean significant difference ($p < 0.05$). See main text for details.

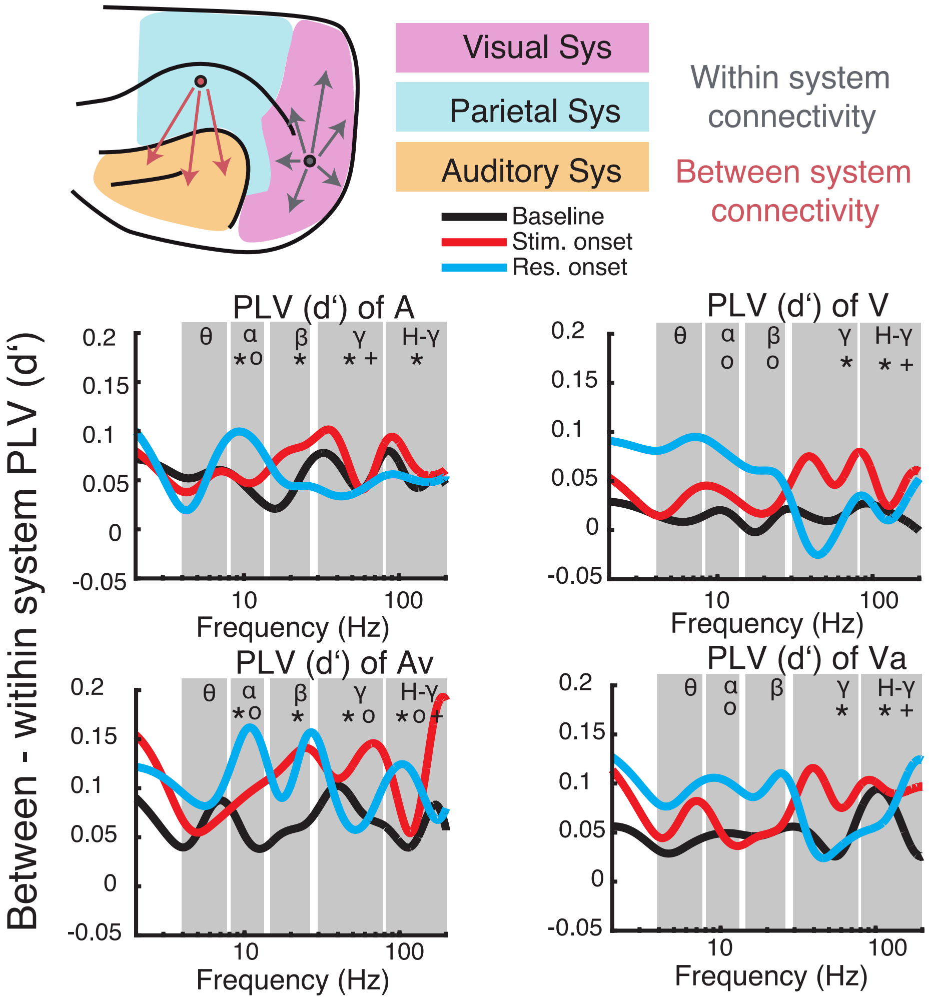


Figure S6. Differences in PLV (in d') between long-range and short-range scales (between and within systems, respectively).
